# Potassium modulates Phytophthora zoospore motility and flagellar beating through dose-dependent effects

**DOI:** 10.64898/2026.10.07.757291

**Authors:** C.A. Lupatelli, A. Attard, E. Galiana, C. Cohen, P. Thomen, X. Noblin

**Author notes:** Co-senior authors and Reference authors: Philippe Thomen.

## Abstract

Biflagellate zoospores are critical agents in the spread of Phytophthora spp., plant pathogens causing major agronomic and economic losses worldwide. These motile cells navigate soil water films toward plant roots using taxis triggered by environmental cues. Here, we investigate how potassium modulates zoospore motility by analyzing trajectory and flagellar beating parameters across KCl concentrations. Using high-speed imaging, we apply kymograph analysis and Fourier transform of the cell body direction over time to quantify flagellar dynamics, while low-speed imaging provides trajectory characteristics. Our results demonstrate that increasing potassium concentration significantly reduces swimming speed while enhancing directional changes. We establish a direct correlation between altered swimming speed and changes in flagellar beating parameters. Our findings indicate that potassium differentially affects anterior and posterior flagella beating. Furthermore, potassium exerts multiple effects, influencing beat amplitude, wave propagation speed and direction-change frequency. This reveals potassium as a key modulator of zoospore motility, triggering a shift from exploratory swimming to confined, reoriented movement. By uncovering the biomechanical responses of zoospores to potassium, this study advances our understanding of how surrounding signals regulate pathogen dispersal in the environment.

## 1 Introduction

*Phytophthora* is a phytopathogenic Oomycete, a genus of eukaryotic filamentous microorganisms classified in the Kingdom of Stramenopiles, alongside heterokont algae like diatoms and brown algae [1]. Among its group, the genus *Phytophthora* counts over 100 species, with a majority causing significant damage to natural and agricultural ecosystems, with economic impacts amounting to billions of dollars annually [2] [3]. A plant infection by *Phytophthora* begins when asexual multinucleate sporangia cleavage and release unicellular, wall-less, bi-flagellated zoospores [4]. These zoospores, capable of moving up to 250 µm/s, spread through water films, droplets, or moist soils, storing energy in the form of lipids necessary to swim for hours [5]. Upon reaching plant surface, they transform into germinative cysts, penetrate host tissue, grow as hyphae and continue the infection cycle which ultimately will produce new zoospores [6]. In this study, we focus on *Phytophthora parasitica*, a soil-borne plant pathogen causing diseases, mainly as rottenness agents, on herbaceous and woody dicotyledonous plants [7].

Zoospores of *P. parasitica* spread disease between roots of neighbouring plants, by swimming through highly complex environments such as soil water film which present strong shear stress zones. The power generated by the two flagella enables them to challenge the viscous forces dominating such small-scaled environment [8]. Morphologically, the zoospores present a kidney-like cell body usually of 10 microns where an anterior flagellum, adorned with tripartite tubular hairs, named mastigonemes, and a posterior smooth flagellum are inserted [4]. The two flagella generate a coordinated thrust by beating together along the anterior-posterior axis, allowing the cell to swim at high velocities during straight runs (250 µm/s) [9]. During turns, a coordinated process resembling a eukaryotic version of the bacterial “run-and-tumble” occurs, where the posterior flagellum halts while the anterior flagellum continues beating, causing the cell body to rotate [9]. This differs for exmaple from the biflagellated *Chlamydomonas* algae which have two identical flagella, attached on the same side that beat synchronously for straight runs and asynchronously for reorientation [10]. A conserved flagellar axonemal structure, composed of 9 doublets of microtubules bound together by dyneins and radial spokes arms, and a motor protein repertoire largely recurrent in other eukaryotic cells, enables the beating orchestration [11].

Zoospores are known for their ability to explore micro-scale environments by exhibiting tactic behaviours [5] and thus constitute an interesting microswimmer model. These biflagellate cells respond to various environmental cues, directing their movement toward the roots of host plants. They display positive rheotaxis and negative geotaxis, enabling them to swim against currents and stay near the soil surface [12] [13]. Additionally, electrotaxis, chemotaxis, and ion currents at the plant root level guide their movement, with zoospores of species like *P. sojae* and *P. cinnamomi* being particularly attracted to plant host exudates [14][15][16][17]. Potassium serves as a crucial signal for zoospores as they navigate water films in the soil. Within this environment, potassium is released due to plant-soil ion exchange dynamics, the activity in the rhizosphere, or the excretion from telluric microorganisms [18]. Previous studies reported how potassium ions modulate zoospores’ swimming and aggregation patterns. For *Phytophthora palmivora*, increased extra-cellular potassium levels reduced swimming speeds and altered movement patterns, such as causing zoospores to swim in circles or in a jerky manner [19]. Depending on the spatiotemporal distributions of potassium levels, *P. parasitica* zoospores may exhibit different motion responses. In areas with higher potassium concentrations, they show a negative chemotactic response, while in other regions, with a specific threshold of 1-4 mM of potassium, they swarm and eventually aggregate [20] [21]. Also changes in intracellular potassium level impact zoospores motion, with some ionophores including gramicidin significantly slowing the swimming speeds of *P. palmivora* and *P. infestans* and changing their swimming patterns from straight to spiral or circular movements [19]. These findings underscore the role of potassium in the cellular mechanisms that control the zoospores’ flagella beating orchestration and ultimately their motion response.

In this article, we focus on quantifying zoospores’ motion response to potassium signals, at different homogenous concentrations. Previous systematic studies on the swimming patterns and spreading characteristics of *P. parasitica* zoospores in water have enhanced our understanding of their biomechanics properties. Kymograph analysis have been previously introduced to investigate zoospores’ flagellar beating [9]. Additionally, new models have been proposed for the hydrodynamic properties of mastigonemes, which are meant to increase forward thrust and to generate additional drag that prevents zoospores from experiencing reversal thrust [9] [22]. Using these tools along with other biophysical approaches new for this Stramenopiles group, we initially examine the overall patterns of zoospore movement in response to potassium, followed by an in-depth analysis of flagella beating patterns.

For the first time, we combine kymograph-based measurements of flagellar beating with Fast Fourier Transform (FFT) analysis to quantify yaw oscillations (rotations about the vertical axis perpendicular to the swimming plane) in *P. parasitica* zoospores. This integrated approach enables us to resolve the beating frequencies of the anterior and posterior flagella with high precision.

Previously, a similar method was used to analyze flagellar beating dynamics in *Chlamydomonas* green algae, where FFT was applied to cell-body oscillations to estimate the average beating frequency across cell populations [23]. More recently, in *Phytophthora infestans* zoospores, FFT was employed to calculate the power spectral density of swimming trajectories, revealing dominant beating frequencies from translational velocity data [24].

In the present study, we extend this framework by investigating, for the first time, cell-body yaw rotations—that is, oscillations about the axis perpendicular to the swimming plane—and their relationship with flagellar beating patterns.

Our observations reveal that as KCl concentration increases zoospores progressively lose the ability to perform long straight swimming runs, and instead show frequent reorientation resulting in high-curved trajectories. These behavior changes were further correlated to flagellar beating. At concentrations below 2 mM KCl, zoospores remain motile but exhibit adaptive changes in their swimming behavior, adjusting both the amplitude and frequency of flagellar beating to maintain propulsion.With higher concentrations of KCl, zoospores’ velocity decreases, as well as the hydrodynamic thrust generated by the two flagella. We discovered that, the eukaryotic “run and tumble” fashion of zoospores is blocked, with the anterior flagellum being the only one to steer the cell body in continuous rotation. Our study elucidates the zoospore response to potassium, offering deeper insights into how zoospores adapt their movement in response to environmental conditions.

### 2 Material and methods

### *2*.*1 Phytophthora* culture conditions and zoospores release

The mycelium of *P. parasitica* (isolate 310, Phytophthora INRA collection, Sophia-Antipolis, France) was routinely cultured on malt agar at 24°C in the dark [18]. For zoospore production, the mycelium was grown for one week in V8 liquid medium at 24°C under continuous light. The mycelium was later drained, macerated, and incubated for an additional five days on 2% water agar to induce sporangia production. Zoospore release was achieved through a heat shock treatment which included incubation of mycelium at 4°C for 30 minutes, followed by the addition of 10 mL distilled water and incubation at 37°C for 20 minutes.

### 2.2 Microscopy observations

We analysed zoospores swimming behaviour in response to discrete concentrations of KCl ranging from 0 to 5 mM. Freshly released zoospores were first mixed with diluted potassium solutions and later injected into a crafted PDMS microfluidic channel (50 or 100 µm height, 1 mm width), presenting an inlet, outlet and a main chamber. We observe the swimming of individual zoospores inside the microchannel under a bright field transmission microscope (Nikon Eclipse Ti2, Minato, Tokyo, Japan) at 40x objective and between 2000 and 4000 fps with the high-speed camera Phantom v711 or v7.3 (Vision Research, NJ, USA). To observe the swimming trajectories of zoospores, we utilized 10x objectives and a frame interval of 0.10 seconds (XIMEA MQ042MG-CM, CO, USA), allowing us to capture larger swimming areas at a lower acquisition speed. The zoospores’ positions were semi-automatically tracked with the Trackmate plugin from Fiji [25] [26].

### 2.3 Determination of trajectory parameters

For the definition of zoospore trajectories, the captured images were processed by Fiji with the Trackmate plugin [27]. To highlight the zoospores from the stacks we first performed a Z-project with average intensity as projection type. We then subtracted from the original stack, the averaged obtained stack and proceeded to make this new stack binary. Trackmate processing criteria were set as follows: estimated blob diameter (LoG detector), 15 µm; Auto initial thresholding; linking max distance (LAP tracker), 30 µm. Only trajectories defined continuously for at least 10 seconds were retained for further treatment. The Trackmate generated data were then treated with a homemade Matlab program to generate velocity histograms, for the calculation of frequency of direction change and of trajectory curvature.

#### Instantaneous speed

It is calculated as following:

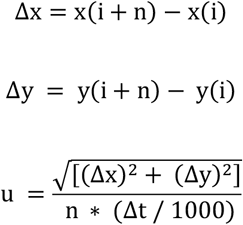

where Δ*x* correspond to x-coordinates between two-time intervals *t*_i_ and *t*_i+n,_ Δ*y* correspond to *y*-coordinates two time intervals *t*_i_ and *t*_(i+n)_. *Δt* is the time interval between two acquired picture and *n* is the averaging factor, here set equal to 2.

#### Frequency of direction change

Displacement vectors were computed using the changes in x and y coordinates (*Δx* and *Δy*). The turning angle *α* between successive displacement vectors over *n* time steps was calculated using the cross product and dot product of the displacement vectors. Local maxima in the *α* vector, where the prominence exceeded 30 degrees, were identified as turning points. The global density of turns by summing the total number of turns across all trajectories divided by the total observation time, provides a comprehensive measure of the zoospores’ turning behavior.

#### Curvature of trajectory

For each trajectory, the x and y coordinates were extracted. Only trajectories longer than 100 µm were selected for curvature analysis. The local radius of curvature (r) was computed at each point along the trajectory using a geometric method based on the circumcenter of three consecutive points. Specifically, for each point *i*, the radius of curvature was derived from the circumscribed circle passing through the triplet of points (*i* − 1, *i, i* + 1). The curvature (curv) was then defined as the reciprocal of the radius (*cutv* = 1/*r*). To avoid edge effects, the curvature values at the first and last points of each trajectory were excluded. The average curvature was subsequently calculated for each trajectory by averaging the remaining curvature values.

### 2.4 Frequency, amplitude and wavelength of flagella beating

We recorded the movement of zoospores during their runs with visible flagella using brightfield microscopy at 40x magnification and a high-speed camera capturing at 2000 or 4000 fps. To analyse flagella beating at discrete concentrations of potassium we proceeded with two separate approaches: by kymograph analysis and by Fourier transform analysis. When flagella were in focus, we determined their beating frequency (*f*), amplitude (*A*) and wavelength (*λ*) as detailed by Tran *et al*. 2022 [9]. Briefly, we applied kymographs to multiple cross-sections perpendicular to the moving direction of zoospores, after stabilizing their position using the Fiji StackReg plugin where possible. The kymographs converted the time-dependent 2D image data into separate signals for analysis [9]. Beating amplitude was quantified from the kymographs by measuring the peak-to-trough distance of individual waveforms and dividing by two, providing the maximal lateral displacement of the flagellum relative to its central axis.

Wavelength was quantified from kymographs drawn parallel to the swimming direction of single swimming zoospores and intersecting each flagellum as shown in Tran et al., 2022 [9]. In the kymographs, flagellar bending waves manifested as oblique, spatially periodic traces. The wavelength λ was defined as the spatial period of the bending wave along the flagellar centerline and was obtained by measuring the distance between two successive points (crest– to–crest) along the spatial axis, and converting this distance from pixels to micrometers.

#### Fourier transform

This method involves using Fourier transform to extract the flagella beating frequency by analysing the oscillation of the cell body angle *versus* time, based on the assumption that the flagella beating wave is reflected in the cell body angular oscillation. This assumption has been used in [23] by the authors, discussed by them as reflecting the fact that forward and backward motion are induced by the beating of flagella which makes the swimmers velocity to fluctuate at that frequency. In the present study, a whole theoretical model is presented in the supplementary section [Supplementary Information S1] to show how the flagellar beating induce angular oscillations of the cell body and justified the data obtained by our new following method. First, we measure such vibration of each zoospore cell body by analyzing their oscillations angle fluctuations (*Ψ*) over time (Figure 1A, Figure 1B). This is done by processing input binary frames using the Analyze Particles tool in Fiji. We then determine the mean angle within the considered frame interval and detrend the angle data by subtracting this mean value from the angle data points within the interval. We apply the Fast Fourier Transform (FFT) algorithm to the detrended angle data, which now represents the amplitude of frequency components in the data. Finally, we store the values in the range of frequencies. The two major peaks identified in the spectrum correspond to the frequencies associated with the two flagella beating (Figure 1C). Due to the conservation of kinetic momentum, the force exerted by each flagellum generates a reaction torque on the cell body, causing it to rotate on itself. Previous studies have shown that the flagella of zoospores propel them in sinusoidal shapes with wave propagation in opposite directions, causing the cell body to gyrate [9]. This asymmetrical beating also causes the cell body to oscillate. By decomposing these yaw vibrational oscillations, we can use FFT to retrieve the components of the propulsion waves from the two flagella. To validate this second method, we also applied it to the previous frames analyzed with kymographs for comparison. For each experiment, we obtained comparable results for the beating frequencies of both the anterior and posterior flagella using kymograph and FFT analysis (Supplementary Information, Figure S2 and Figure S3). The difference between the two methods when analyzing the same frames was around a maximum of 15%. This indicates that both methods are reliable and can be used effectively for determining flagella beating frequencies.

**Figure 1.**
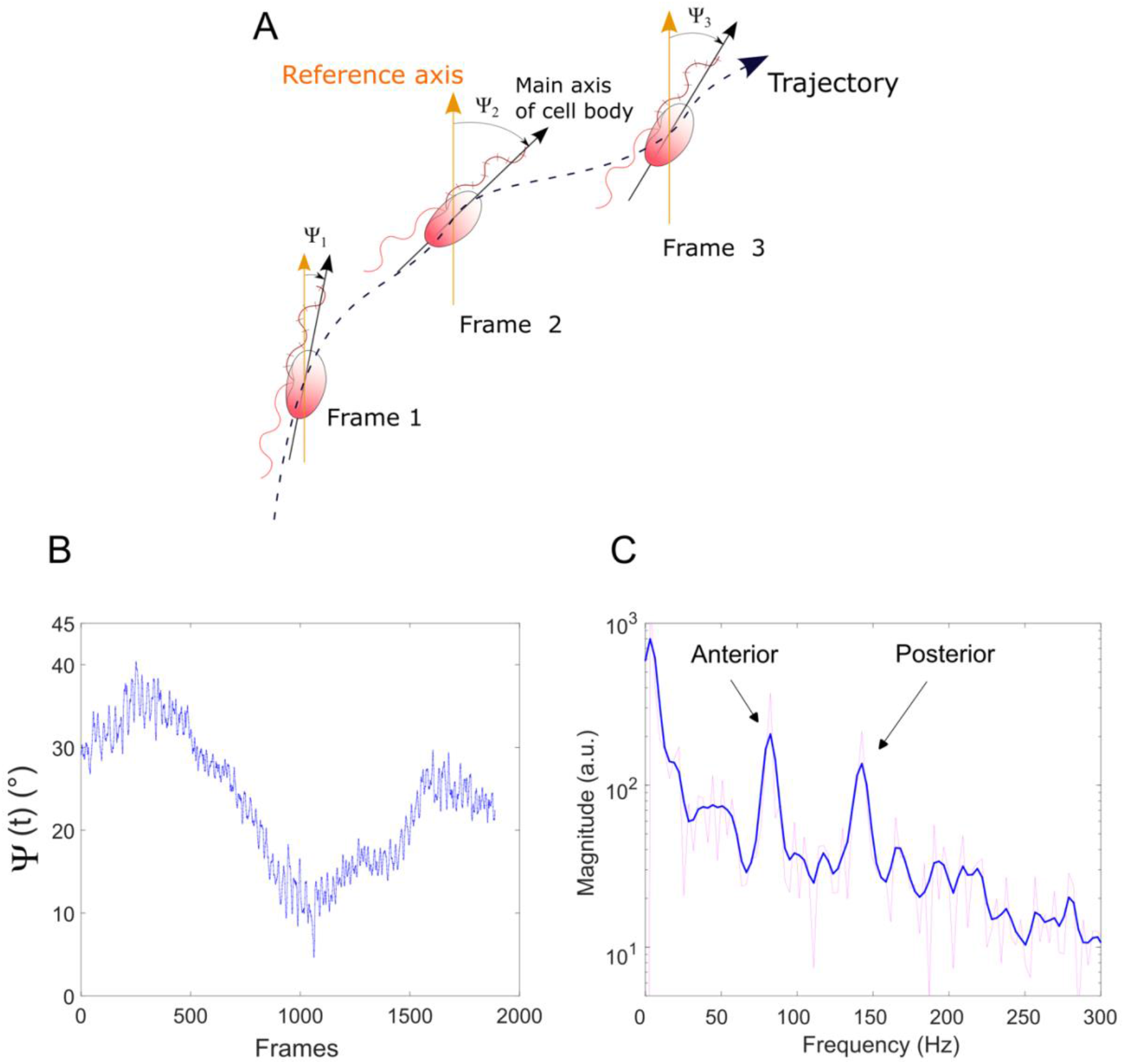
Fourier Transform analysis of zoospore flagellar beating. (A) The initial step involves extracting the oscillation angle (Ψ) of the zoospore cell body as a function of time, measured from the body’s oscillation motion. (B) The graph shows the oscillation of the cell body angle Ψ over time (expressed in frames). (C) The Fourier Transform of this oscillatory motion reveals two distinct peaks corresponding to the anterior and posterior flagellar beat frequencies. Blue curve is a 5-points smoothing of the spectrum and the red curve is the raw spectrum. In this example, measured at 1.5 mM, the first peak represents the anterior flagellum at approximately 82 Hz, and the second peak corresponds to the posterior flagellum at around 143 Hz.

Statistical analysis and graphic representations were performed using GraphPad Prism version 10 (GraphPad Software, Boston, Massachusetts USA).

## 3 Results

### 3.1 Impact of potassium on zoospores’ kinematics

We investigate the characteristics of zoospore swimming by analyzing their trajectories and behaviours at increasing concentrations of KCl. To perform microscopic assays, we first subject zoospores to a KCl solution with different concentrations ranging from 0 to 5 mM. We then confined them in simple manufactured microfluidic devices to maximize the likelihood of keeping them in focus during observations. The swimming of zoospores was recorded at 10x magnification with a frame interval of 0.10 s for 10 to 40 seconds.

Figure 2 shows typical trajectories for concentrations of potassium ranging from 0 to 5 mM. At low concentrations of potassium (0-1 mM), zoospores perform the typical long and straight runs separated by few random turning events, in a run-and-tumble fashion as reported previously [9] [28]; their mean speed is around 200 µm/s (197 ± 36 µm/s at 0 mM; 172 ± 48 µm/s at 0.5 mM; 209 ± 44 µm/s at 1 mM). At 1.5 and 2 mM potassium concentration, the trajectories of the zoospores remain straight, but they are more likely to alternate between direction changes and straight runs; their mean speed is reduced (136 ± 55 µm/s at 1.5 mM; 128 ± 65 at 2 mM). At 2.5 and 3 mM, zoospore trajectories exhibit an increasing number of turns, resulting in smoother and more curved paths compared to the well-defined, straight trajectories observed between 0 and 1 mM KCl. It is combined with a decrease in mean speed (87 ± 44 µm/s at 2.5 mM). At 5 mM, the zoospores swim in a spiral fashion with small concentric curves, limiting their displacement from the initial point and indicating that they are likely turning almost on the same spot. From 3 to 5mM, the mean speed goes on to decrease (28 ± 18 µm/s at 3 mM; 12 ± 7 µm/s at 5 mM).

**Figure 2.**
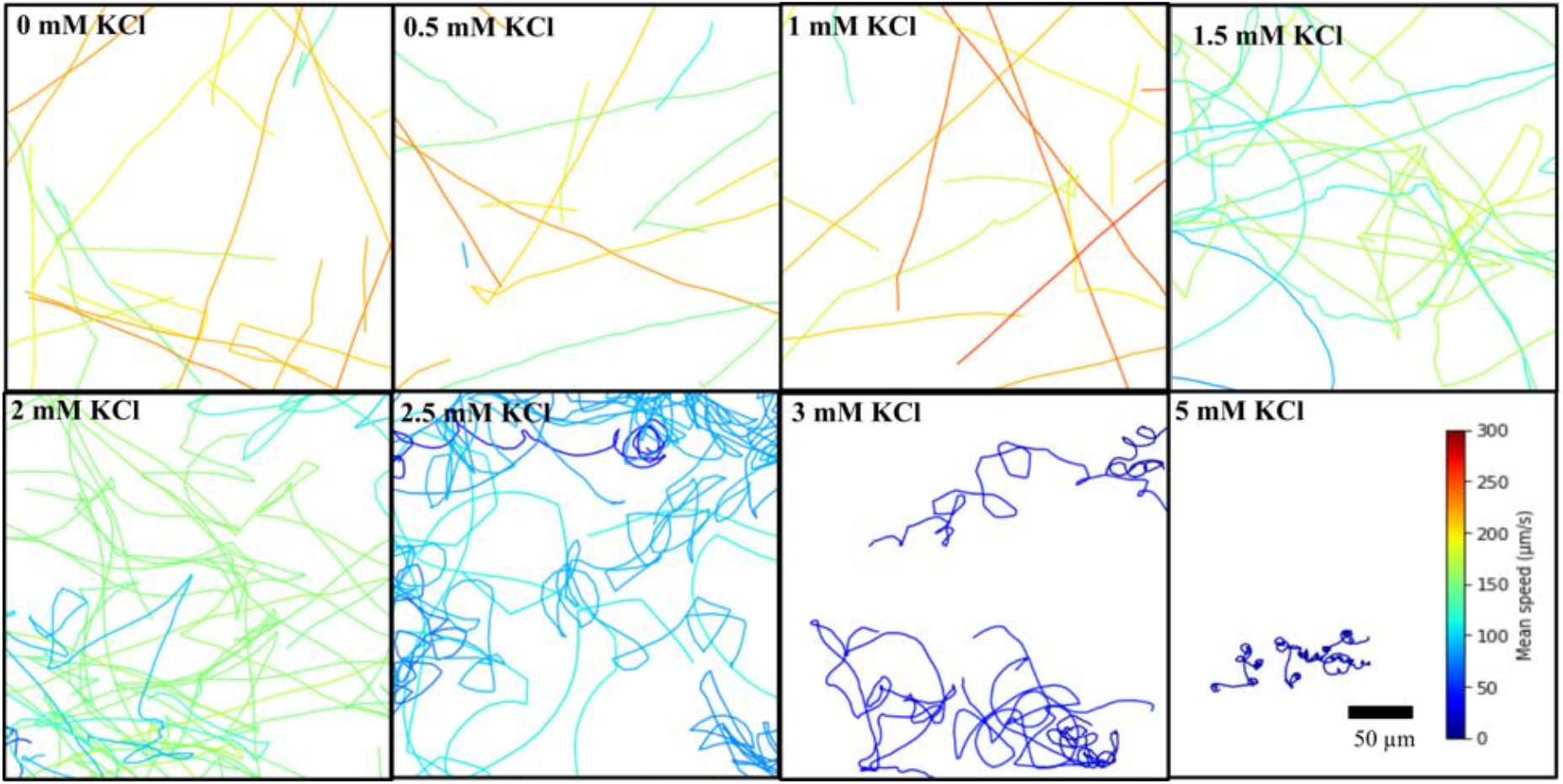
Zoospore trajectories as a function of KCl concentration. At 0, 0.5 and 1 mM KCl, zoospores predominantly exhibit straight-line trajectories with occasional turns. At 1.5 and 2 mM KCl, there is an increase in the frequency of sharp turns along their paths and the mean speed is reduced. From 2.5 mM KCl onward, the zoospore trajectories become increasingly erratic, characterized by more frequent turns and circular patterns, with a decreasing speed. The color indicates the mean speed of each trajectory according to the colorbar.

To be more quantitative, we estimated instantaneous speed distribution *p(U)* of each zoospore, using their *x* and *y* position data, at increasing potassium concentrations (Figure 3). The speed distribution *p(U)* presents in normal conditions (low potassium) a combination of two different normal distributions which has been interpreted as two behavioral states of running and turning [9]:

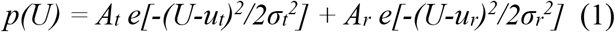

where *u*_*t*_ and *u*_*r*_ are the mean speed for the turning and running events respectively. We proposed to assume such behavior for any potassium concentration and try to fit the data in that way. Each distribution from 0 mM to 2 mM KCl can be fitted to obtain the values of *u*_*t*_ and *u*_*r*_. The results show that the occurrence of turning events increases notably at 1.5 mM and 2 mM KCl, corroborating the qualitative observations made on trajectories, and responsible for the reduced mean speed of the zoospores. Between 0 and 1 mM of KCl, mean values of *U* are around 200 µm/s; above 1mM potassium, the range of instantaneous speed begins to drop, with mean value of *U* decreasing from 136 µm/s at 1.5 mM KCl to 128 µm/s at 2 mM. Interestingly, both running and turning event speeds take their highest value at 1 mM KCl.

**Figure 3.**
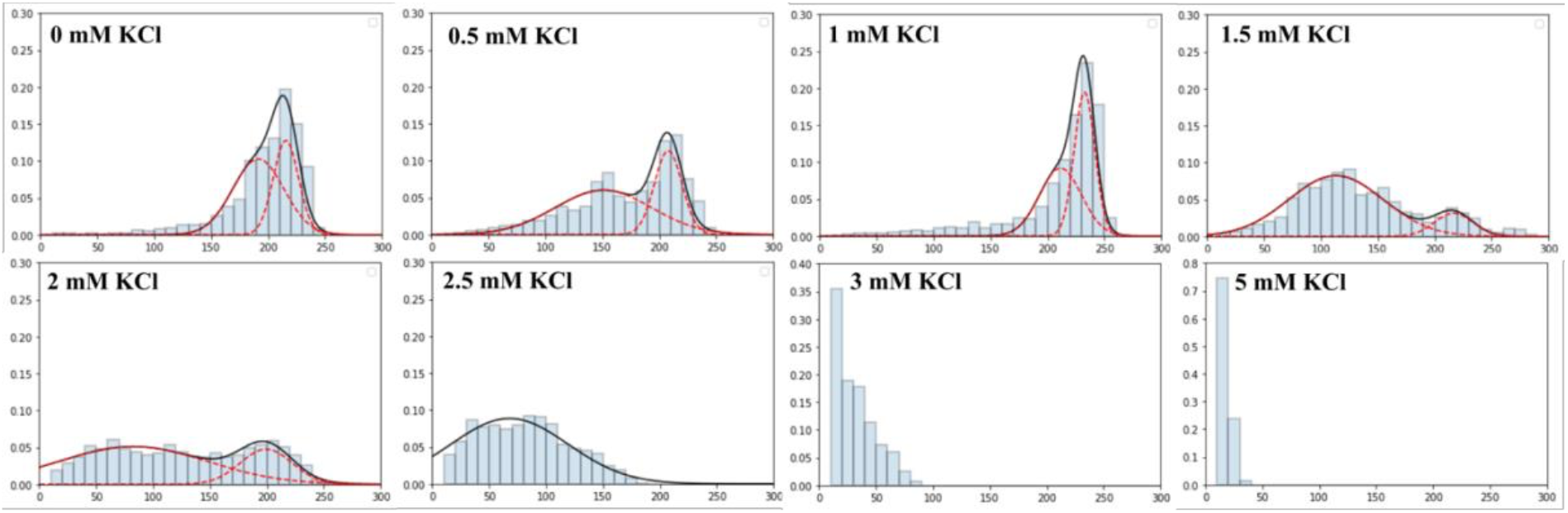
Instantaneous speed distribution of zoospores in response to varying KCl concentrations (0–5 mM). Zoospores exhibit high swimming velocities, with a pick above 200 µm/s from 0 to 1 mM KCl. At 1.5 and 2 mM KCl, the speed distribution becomes broader, with a large distribution of low velocities peak around 100 µm/s. At 2.5 mM, the instantaneous speed still declines, with most values ranging between 50–100 µm/s. At 3 mM and 5 mM KCl, the majority of velocities are distributed below 80 µm/s and 30 µm/s, respectively. From 0 to 2 mM, the histograms were fitted with the two Gaussians of equation (1) (red doted lines for the Gaussians and black line for the sum). At 2.5 mM, data were fitted with one Gaussian.

From 2.5 mM KCl, the speed distribution is no longer fitted by two Gaussians, in accordance with the qualitative observations of the trajectories that no longer show sequences of long straight lines and turns. The mean speed drops to 87 µm/s at 2.5 mM KCl; at 3 mM zoospores mostly swim below 80 µm/s and at 5 mM they almost stop moving with instantaneous speed below 30 µm/s.

To further analyze the zoospores’ swimming patterns in response to potassium, we estimated the total number of turning events per second (Figure 4A) and the curvature of their trajectories (Figure 4B). We defined a single turning event as an instance where the angle *α* between two successive displacement vectors exceeded a local prominence of 30°, where each displacement vector was computed over an interval of n = 3 consecutive frames (corresponding to 0.30–0.36 s, depending on the acquisition frame rate; inter-frame interval Δt = 0.10 or 0.12 s across movies). Turning events were therefore detected over a total time window of 0.60–0.72 s (2 × 3 frames), and only local angle maxima meeting the 30° prominence threshold were retained to avoid counting minor fluctuations in trajectory direction.

**Figure 4.**
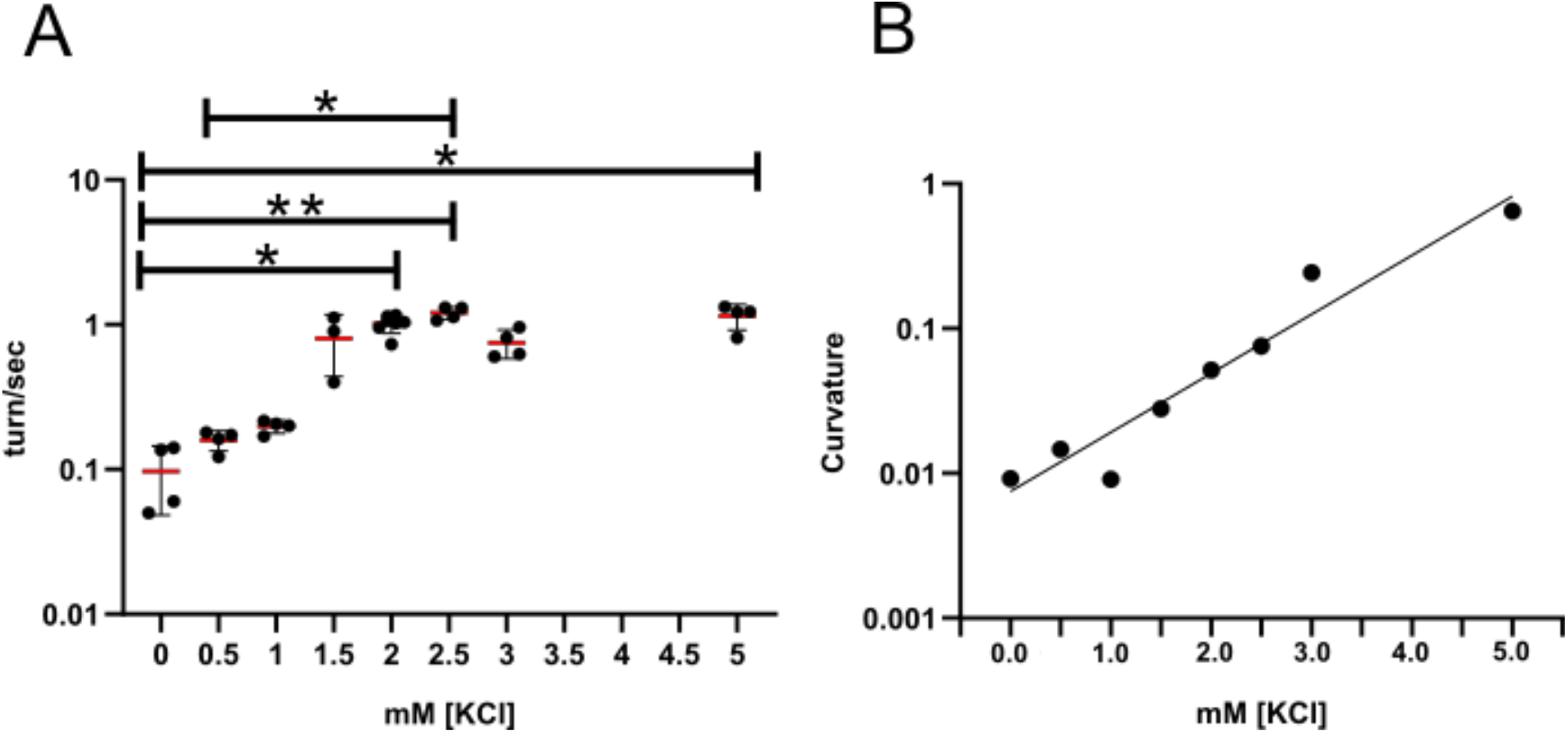
Characteristics of zoospore trajectories as a function of increasing KCl concentrations. (A) The number of turns in zoospore trajectories per second. From 1.5 mM KCl, zoospores exhibit an average of one turn per second. Significant statistical differences were observed using the Kruskal-Wallis test between 0 and 2 mM (p = 0.0350), 0 and 2.5 mM (p = 0.0054), 0 and 5 mM (p = 0.01), and 0.5 and 2.5 mM (p = 0.0345). (B) The curvature of zoospore trajectories (dots) as a function of potassium concentration. Doted line is an exponential fit of the data with the function y([KCl]) = A exp([KCl]/C_0_) with A = 0.0075 and C_0_ = 1.065 mM.

Accordingly, at 0 mM KCl, we observe that zoospores swim with 0.1 turning events per second; the rate of turns increases when potassium is increased from 0 to 1.5 mM, up to 1 turning event per second. Beyond 1.5 mM KCl,a plateau-like trend where zoospores encounter approximately one turning event per second, is observed. We then estimated the curvature of zoospore trajectories by calculating the inverse of the radius of curvature computed for each trajectory point, where larger radius values indicate smaller curvatures. Consistently, we found that higher KCl concentrations lead to increased trajectory curvature, reflected by decreasing radius values, and resulting in highly circular paths readily visible in Figure 2. The increase in curvature is well fitted by an exponential function of potassium concentration.

### 3.2 Potassium differentially influences anterior and posterior flagella beating

The swimming pattern is regulated by the beating of the two flagella positioned in opposite directions along the cell body’s anterior-posterior axis [9]. In this section, we show how the beating pattern of each flagellum is affected in presence of potassium and how it modulates zoospores’ speed and trajectories in response to increasing KCl concentrations. This analysis is based on microscopy assays at a small scale, where the flagella are visible and observed over short periods (from 0.25 s to 2 s).

We recorded the movement of zoospores at high speed imaging and high magnification. We observe that the beating frequencies of anterior and posterior flagella are modulated differently with increasing concentrations of KCl (Figure 5A-5B). The beating frequency of the anterior flagellum increases from 0 to 1 mM KCl, reaching an average of 98 Hz, before progressively decreasing at 1.5 and 2 mM to values below 80 Hz. At 2.5 mM, frequencies drop below 50 Hz. At 3 and 5 mM KCl, FFT analysis yielded only sparse peaks, suggesting beating below 35 Hz, while kymograph analysis was not feasible. At these higher concentrations, as confirmed by trajectory analysis, zoospores move in persistent circular paths, complicating the acquisition of frames perpendicular to the anterior flagellum axis. The beating frequency of the posterior flagellum remains relatively stable between 0 and 2 mM KCl, averaging around 140 Hz. At 2.5 and 3 mM KCl, only limited FFT measurements were feasible, and kymograph analysis could not be performed, due to the same constraints outlined above. At 5 mM KCl, no frequency could be measured; as shown in the supplementary video, the posterior flagellum appeared to cease beating entirely

**Figure 5.**
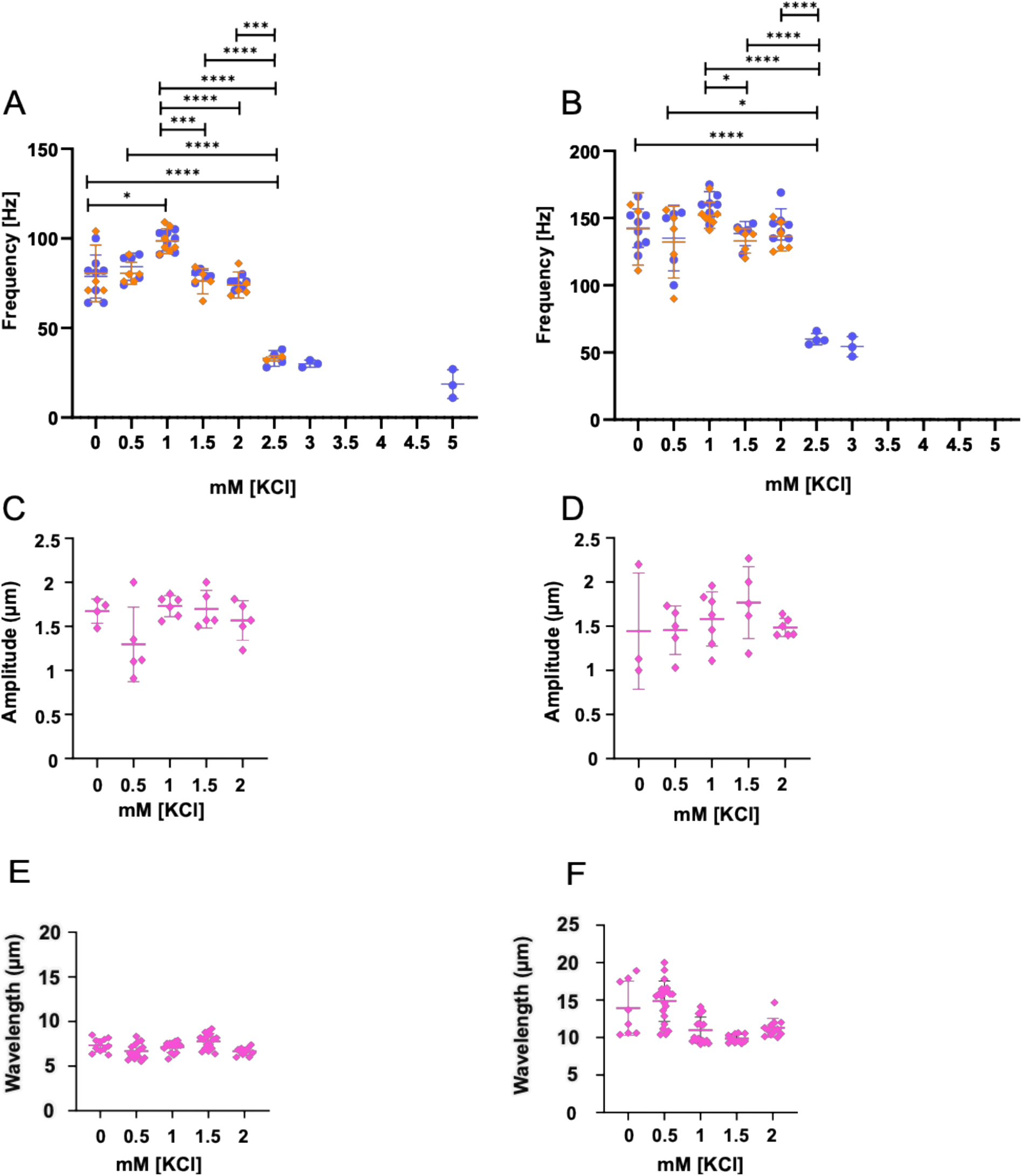
Analysis of flagellar beating parameters at increasing KCl concentrations. Beating frequencies were obtained using Fourier transform and kymograph analysis: (A) Anterior flagellar beating frequencies (in blue for Fourier transform analysis and orange for kymograph analysis) increases up to 1 mM KCl, followed by a gradual decrease at higher concentrations; Brown-Forsythe and Welch ANOVA tests: adjusted P values are * 0.017, *** <0.0002, **** <0.0001; (B) Posterior flagellar beating frequencies (in blue for Fourier transform analysis and orange for kymograph analysis), show a plateau up to 2 mM KCl, followed by a sharp decline, eventually stopping completely at 5 mM KCl; Brown-Forsythe and Welch ANOVA tests: adjusted P values are * <0.026, **** <0.0001. Beating amplitudes were obtained using kymograph: Neither the beating amplitude of the anterior flagella (C) nor the beating amplitude of the posterior flagellar (D) vary significantly as a function of potassium. As amplitude is deduced by kymographs, we were able to get amplitude only up to 2 mM KCl. Wavelengths were obtained using kymograph: The wavelngth of the anterior flagella does not vary as a function of potassium concentration (E); the wavelngth of the posterior flagella show a decrease when the potassium is increased from 0.5 to 1 mM (F). Each point is a measurement of one of the above parameters on one flagella (anterior or posterior) of a one zoospore, in one replicate.

Kymograph analysis revealed no significant changes in the beating amplitude of either the anterior or posterior flagellum as potassium concentration increase from 0 to 2 mM (Figure 5C-D). No effect on wavelength of the aneterior flagellum is observed, but a diminution of about 25% is observed in wavelength of the posterorior flagellum at 1 mM and above compared to smallest potassium concentrations (Figure 5E-F).

## 4 Discussion

*P. parasitica* zoospores are highly proficient microswimmers, capable of moving through thin water films and propagating disease between the roots of neighbouring plants in soil. Previous studies have reported that *Phytophthora* zoospores exhibit a tactic response to potassium, which can vary based on its spatiotemporal distribution [19] [20]. While the molecular mechanisms of potassium, or more in general signal sensing and cellular response regulation in zoospores are not yet fully understood, some components of their sensory membrane repertoire have started to be investigated [13] [29] [30]. Here, we present novel quantifications of this chemotactic response, providing new insights into the regulation of zoospore motion in the presence of external signals.

From low-speed imaging, three regimes can be distinguished. In a first regime, from 0 to 1 mM KCl, zoospores of *P. parasitica* display typical long and straight trajectories with sustained velocities around 200 µm/s, exhibiting a typical eukaryotic run-and-tumble behaviour [31]. In a second and transient regime, at 1.5 and 2 mM KCl, two peaks are still observed in the histogramm of instantaneous speed, but with a large occurrence of low speed; the frequency of direction changes reach a maximum. In fact, the frequency of direction changes increases exponentially with time, from 0.1 to 1 change per second when potassium is increased from 0 to 2 mM. In a third regime, from 2.5 mM KCl onward, zoospore velocity sharply decreases and no peak at 200 µm/s are observed in the histogramm of instantaneous speed. The trajectory curvature increases exponentially with potassium concentration. This phenotype is consistent with observations by Appiah *et al*. in *P. palmivora* zoospores, where the addition of KCl reduced swimming speed and, from 4 mM KCl, caused zoospores to swim in perpetual circles [19]. At very high potassium concentrations (3 and 5 mM), the turns exhibited a pronounced curvature resulting in progressively short-circle trajectories. During chemotaxis and phototaxis, other organisms, including sperm, algae, and marine zooplankton, also tend to exhibit drifting circular or altered helical paths [32] [33].

High-speed imaging reveals the responses in the beating frequencies, amplitudes and wavelength of both flagella to increasing KCl concentrations. We used the Fourier transform to measure the beating frequency of the flagella. The linearity of the Stokes equation ensures that the time-dependent variation of the cell body angle arises from additive contributions from each flagellum. In Fourier space, this corresponds to two distinct peaks. However, this method, in its current form, does not provide direct access to the beating amplitude. In future theoretical work, a more detailed quantitative analysis of the peak heights could potentially yield a reliable estimate of the beating amplitude in the future. In the current study, the results obtained from both kymograph and FFT analysis are consistent, demonstrating that FFT can be effectively used to calculate beating frequencies when focused images of flagella cannot be captured, or kymograph analysis on perpendicular flagella sections are not possible.

In the 0–2 mM potassium range, where speed histograms show a peak around 200 µm/s and trajectories display extended straight segments, both flagella maintain a high beating frequency. The beating frequencies of both the anterior and posterior flagella reach their maximum values at 1 mM KCl. At 2.5 mM and above, both flagella exhibit a marked decrease in beating frequency, falling below their baseline values in water. This reduction in flagellar beating frequencies correlates with a drastic decline in the zoospores’ instantaneous swimming speed, as measured by low-speed imaging.

During directional changes, the posterior flagellum of zoospores typically ceases beating entirely, while the anterior flagellum continues to beat in a stroke-like manner until the cell body is oriented in a new direction and resumes straight-line swimming [9]. In our study, however, zoospores exposed to high KCl concentrations (5 mM) failed to transition back to straight-line swimming with both flagella beating. As shown in the Supplementary Videos, they instead remained trapped in continuous circular motion, rotating anticlockwise in place, primarily propelled by the anterior flagellum.

These observations suggest that to resume straight-line trajectories, zoospores may require an environmental stimulus, such as a local decrease in potassium concentration. Under our experimental conditions (homogeneous KCl concentration), this stimulus is absent. In a potassium concentration gradient, zoospores might actively reorient toward lower-concentration zones, as demonstrated earlier in [20], where efficient avoidance of high-potassium areas were observed. Thus, swimming behaviors are expected to differ depending on whether zoospores are exposed to homogeneous or gradient conditions—a hypothesis warranting further investigation in controlled potassium gradients.

Finally, the theoretical model presented in the Supplementary Information predicts that curvilinear trajectories could also emerge when both flagella are beating, provided their beating frequencies are differentially modulated. Although such behaviors were not observed in our study, this prediction highlights the critical role of inter-flagellar coordination dynamics in trajectory regulation.

The beating frequencies of both the anterior and posterior flagella reach their maximum values at 1 mM KCl. At 2.5 mM and above, both flagella exhibit a marked decrease in beating frequency, falling below their baseline values in water. This reduction in flagellar beating frequencies correlates with a drastic decline in the zoospores’ instantaneous swimming speed, as measured by low-speed imaging.

During directional changes, the posterior flagellum of zoospores typically ceases beating entirely, while the anterior flagellum continues to beat in a stroke-like manner until the cell body is oriented in a new direction and resumes straight-line swimming [9]. In our study, however, zoospores exposed to high KCl concentrations (5 mM) failed to transition back to straight-line swimming with both flagella beating. As shown in the Supplementary Videos, they instead remained trapped in continuous circular motion, rotating anticlockwise in place, primarily propelled by the anterior flagellum.

These observations suggest that to resume straight-line trajectories, zoospores may require an environmental stimulus, such as a local decrease in potassium concentration. Under our experimental conditions (homogeneous KCl concentration), this stimulus is absent. In a potassium concentration gradient, zoospores might actively reorient toward lower-concentration zones, as demonstrated earlier in [20], where efficient avoidance of high-potassium areas were observed. Thus, swimming behaviors are expected to differ depending on whether zoospores are exposed to homogeneous or gradient conditions—a hypothesis warranting further investigation in controlled potassium gradients.

Finally, the theoretical model presented in the Supplementary Information predicts that curvilinear trajectories could also emerge when both flagella are beating, provided their beating frequencies are differentially modulated. Although such behaviors were not observed in our study, this prediction highlights the critical role of inter-flagellar coordination dynamics in trajectory regulation.

Amplitudes of both the anterior and posterior flagella do not vary significantly except for the anterior flagellum at 0.5 mM where a decreased amplitude is observed. The outcome is a reduced speed, as measured both in low and high speed imaging (see below).

The wavelength of anterior flagellum is constant when increasing potassium concentration; wavelength of posterior flagellum exhibits a decrease of about 25%, but as shownbelow, this variation does not contribute to a change in velocity.

To reinforce our observations, we proceeded to the calculation of zoospores speed using the Resistive Force Theory [9]. In the formula the speed depends on beating parameters: amplitude, frequency and wavelength. For this study, we restricted the analysis to potassium concentrations between 0 and 2 mM, a range for which complete single-cell measurements of flagellar beating frequency, beat amplitude, and wavelength were available for individual zoospores. The speed can then be predicted for each zoospores for which all parameters are accessible, but as there are only few zoospores for which all parameters are measured, we also calculated the mean speed expected, using mean values of amplitude, frequency and wavlength, for each KCl concentration. The mean speeds calculated for each concentration are compared with the mean speed measured from low-speed imaging (Figure 6). The same trends are observed, the most striking being the decrease in speed observed at 0.5 mM. The relatively low speed observed at 0.5 mM is mainly explained by a decrease in the beating amplitude of the anterior flagellum. The high speed at 1 mM, in contrast, is mainly explained by a high propagation speed of the beating wave of the anterior flagellum, due to the high value of the beating frequency. It is worth noting that, in line with what was observed earlier [9], zoospore speed is mainly due to the beating of the anterior flagellum.

**Figure 6.**
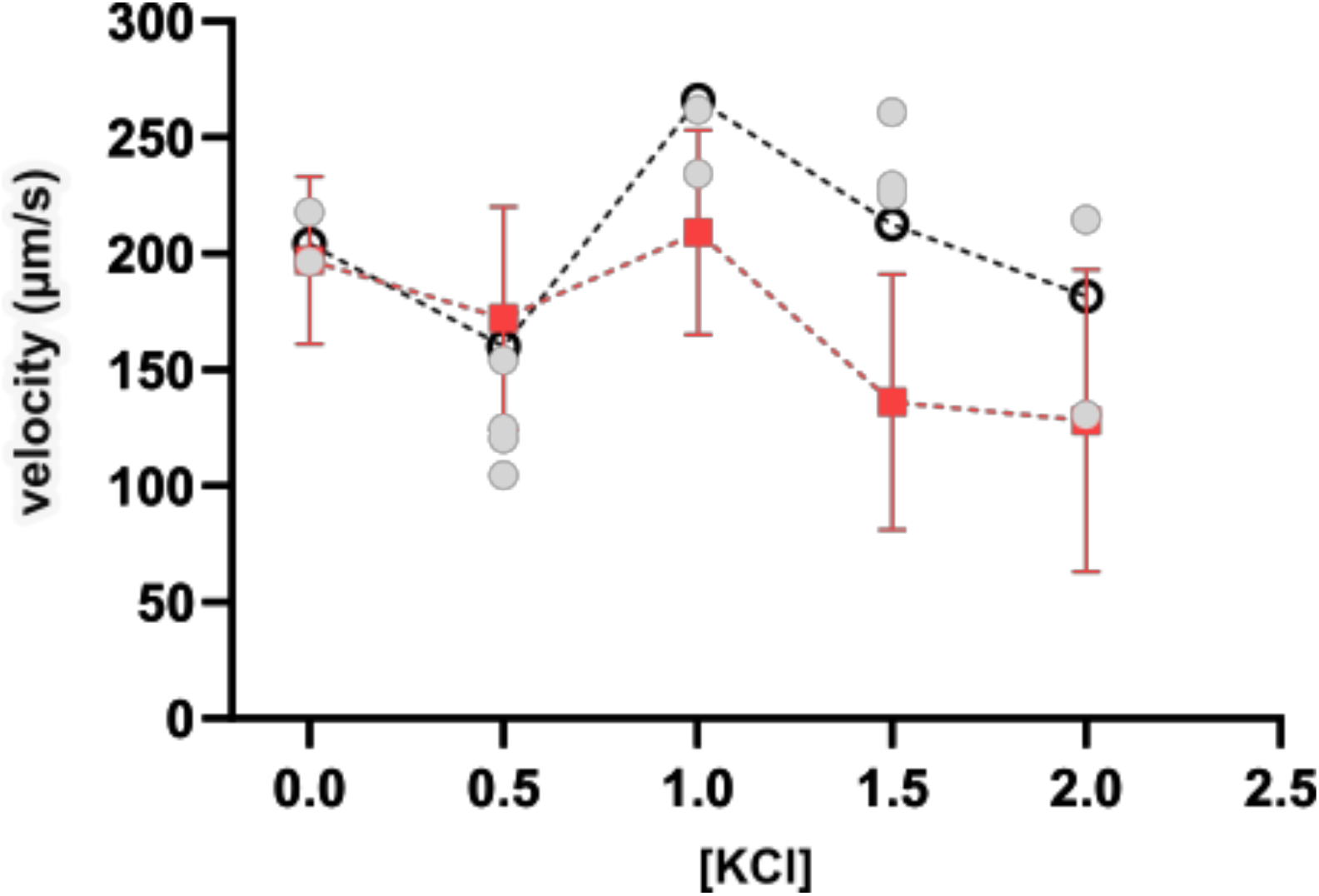
Mean speed of zoospores as a function of potassium concentration. The speed has been calculated using RFT theory (see text) with the beating parameters measured in the current study from high speed imaging (open black circles). The whole parameters were available only on few zoospores for which the speed was calculated with RFT theory (gray circles). The red squares correspond to the mean speed calculated from low speed imaging associated with standard deviation.

Interestingly, as the potassium concentration increases from 0 to 0.5 mM and then to 1 mM, the frequency of direction changes in zoospore swimming increases in an approximately exponential manner. Potassium thus exerts multiple effects on zoospore motility: (i) it influences both the beating amplitude (at 0.5 mM KCl) and the wave propagation speed (the product of beating frequency and wavelength), two key parameters that determine swimming speed; (ii) it also affects the frequency of direction changes, thereby altering the trajectory pattern. However, these effects do not appear to be correlated a priori, although this possibility cannot be ruled out based on the current data.

In Figure 6, the differences at 1.5 and 2 mM between data obtained with low-speed imaging and data obtained with high-speed imaging can be explained by the fact that low-speed imaging accounts for all velocities (straight line portions and turns where low speeds are expected), whereas high-speed imaging computation selects rectilinear portions of trajectories, where the highest velocities are expected to be observed.

Taken together, from an environmental perspective, these results align with the observed negative taxis of zoospores in potassium gradients (1-4 mM) and their subsequent aggregation on artificial surfaces [20]. The increasing velocities up to 1.5 mM of KCl, likely can enable zoospores to avoid high K^+^ concentrations and migrate toward the root surface, where potassium levels are typically lower due to proton uptake [34] [35]. Conversely, a high potassium concentration might inhibit zoospores’ ability to reorient their trajectory and recover normal swimming patterns, with zoospores unable to move long distances, remaining “trapped” within same area. Further investigations using more sophisticated experimental setups with controlled potassium gradient generators are needed. Additionally, complementary biological analyses—such as gene knockouts and motion tracking of zoospores lacking key signal-sensing components, including potassium receptors—would help clarify the underlying mechanisms governing zoospore taxis and adaptation [30]. Based on observations made in other eukaryotic microorganisms, we can propose a mechanistic pathway where elevated extracellular K^+^ causes membrane depolarisation, subsequently activating voltage-gated calcium channels and leading to increased cytoplasmic Ca^2+^ levels that disrupt flagellar coordination through calcium-induced desynchronisation [36]. Recent transcriptomic evidence suggesting involvement of specialised K^+^/Ca^2+^ channels and α-carbonic anhydrase in K^+^ electroception further supports this model, indicating that potassium acts through dedicated cation transport systems that integrate with established chemotactic machinery [29]. Future work should focus on directly measuring membrane potential changes in response to K^+^ gradients, identifying specific K^+^-sensitive receptors, and characterising how calcium oscillations control flagellar beating patterns and directional responses to environmental potassium cues.

## 5 Conclusion

In this study, we quantified the motion response of *P. parasitica* zoospores to increasing concentrations of potassium. Previous systematic studies have enhanced our understanding of the swimming patterns and biomechanics of properties of zoospores in water. Chemotactic responses to potassium have also been documented. Here, we first investigate the overall characteristics of zoospore trajectories in response to potassium, then delve into the swimming mechanisms modulated at the flagella level.

We observed that with increasing KCl concentrations, zoospores are unable to perform long straight runs and instead exhibit frequent directional changes, resulting in highly curved trajectories and decreased instantaneous speeds. This global swimming phenotype resulted associated to abnormal motion regulation by the two flagella. For the first time in *Phytophthora* zoospores, we analyzed flagella beating using both kymograph and Fourier transform analysis based on single cell bodies vibrations. We discovered that the two flagella respond differently to increasing potassium concentrations. The anterior flagellum shows a significant increase and subsequent decrease in beating frequency, while the posterior flagellum’s response is less pronounced. At high potassium concentrations, the anterior flagellum fails to reorient the cell body, and the posterior flagellum completely stops beating at 5 mM, resulting in perpetual circular trajectories driven by the anterior flagellum.

Our study elucidates the mechanisms and characteristics of zoospore response to potassium, providing deeper insights into their motion adaptation in relation to their environment. This work advances our understanding of the intricate dynamics governing zoospore motility and their interaction with external stimuli.

## FIGURES

**Figure S2.**
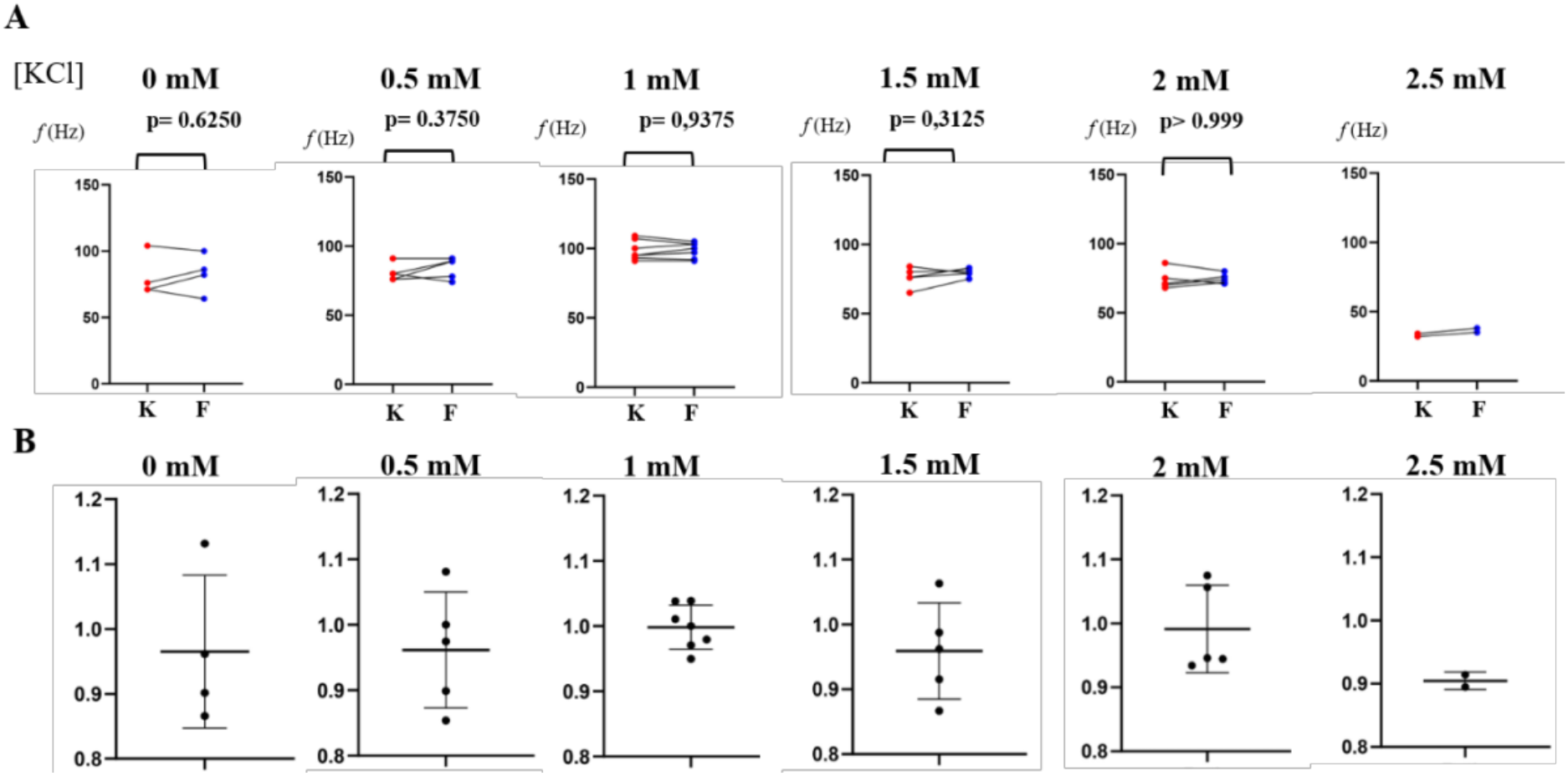
Comparison between kymograph and Fourier Transform analysis for anterior flagellum under increasing potassium concentrations. (A) Correlation between results from the same experiment obtained using kymograph (red) and Fourier transform (blue) analysis. Paired Wilcoxon test revealed no statistically significant differences between the means (p > 0.05), indicating consistent results across both methods. (B) Estimated ratio of kymograph to Fourier Transform results (kymograph/Fourier ratio), with no discrepancies exceeding 15%, further validating the reliability of both analytical approaches.

**Figure S3.**
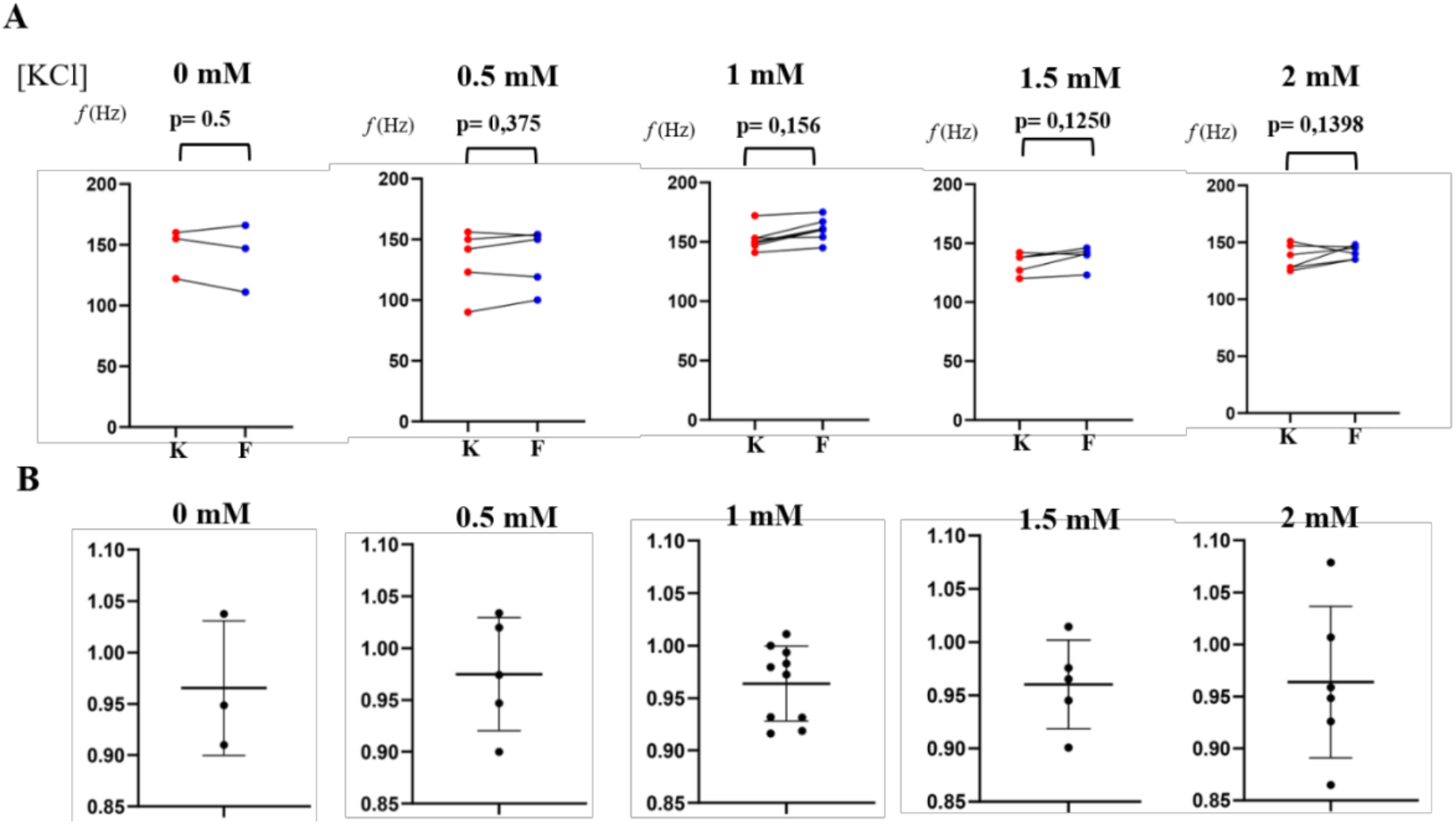
Comparison between kymograph analysis and Fourier Transform for posterior flagellum under increasing potassium concentrations. (A) Correlation between results from the same experiment obtained using kymograph (red) and Fourier Transform (blue) analysis. Paired Wilcoxon test revealed no statistically significant differences between the means (p > 0.05), indicating consistent results across both methods. (B) Estimated ratio of kymograph to Fourier Transform results (kymograph/Fourier ratio), with no discrepancies exceeding 12%, further validating the reliability of both analytical approaches.

**Tables S4.**

| [KCl] | Number of experiments | Number of trajectories | Number of instantaneous speeds |
| --- | --- | --- | --- |
| 0 mM | 4 | 130 | 3027 |
| 0.5 mM | 4 | 151 | 3035 |
| 1 mM | 4 | 174 | 3686 |
| 1.5 mM | 3 | 23 | 1347 |
| 2 mM | 7 | 29 | 3150 |
| 2.5 mM | 4 | 18 | 3840 |
| 3 mM | 4 | 15 | 1532 |
| 5 mM | 4 | 5 | 965 |

Number of experimental repetitions for data used in the calculus of the density of turns (figure 4A):
| [KCl] | Number of replicates | Number of trajectories |
| --- | --- | --- |
| 0 mM | 4 | 33 |
| 0.5 mM | 4 | 45 |
| 1 mM | 4 | 60 |
| 1.5 mM | 3 | 22 |
| 2 mM | 3 | 29 |
| 2.5 mM | 4 | 19 |
| 3 mM | 4 | 7 |
| 5 mM | 4 | 6 |

Number of experimental repetitions for data used in the calculus of the curvature of the trajectories (figure 4B):
| [KCl] | Number of replicates | Number of trajectories |
| --- | --- | --- |
| 0 mM | 4 | 25 |
| 0.5 mM | 4 | 19 |
| 1 mM | 4 | 31 |
| 1.5 mM | 3 | 23 |
| 2 mM | 3 | 25 |
| 2.5 mM | 4 | 19 |
| 3 mM | 4 | 7 |
| 5 mM | 4 | 6 |

## Supplementary Informations

Supplementary material to the manuscript “…”

16 avril 2026

Modeling of cell body angular oscillations during zoospores swimming to explain the FFT spectrum obtained.

## 1 Introduction

We describe here how the two flagella beating influence the angular oscillations of the main direction of the cell body **compared** to a fixed direction. We note Ψ this angle with the horizontal axis in the lab frame (see Fig. S1). We start from the equation of Resistive Force Theory (RFT). Compared to its use in [1], it needs to be adapted as not only the projection on the cell body axis is important. The force balance projected in the direction of motion is **insufficient** and equations relative to kinetic momentum (then cell body rotation) have to be exploited.

**Figure S1:**
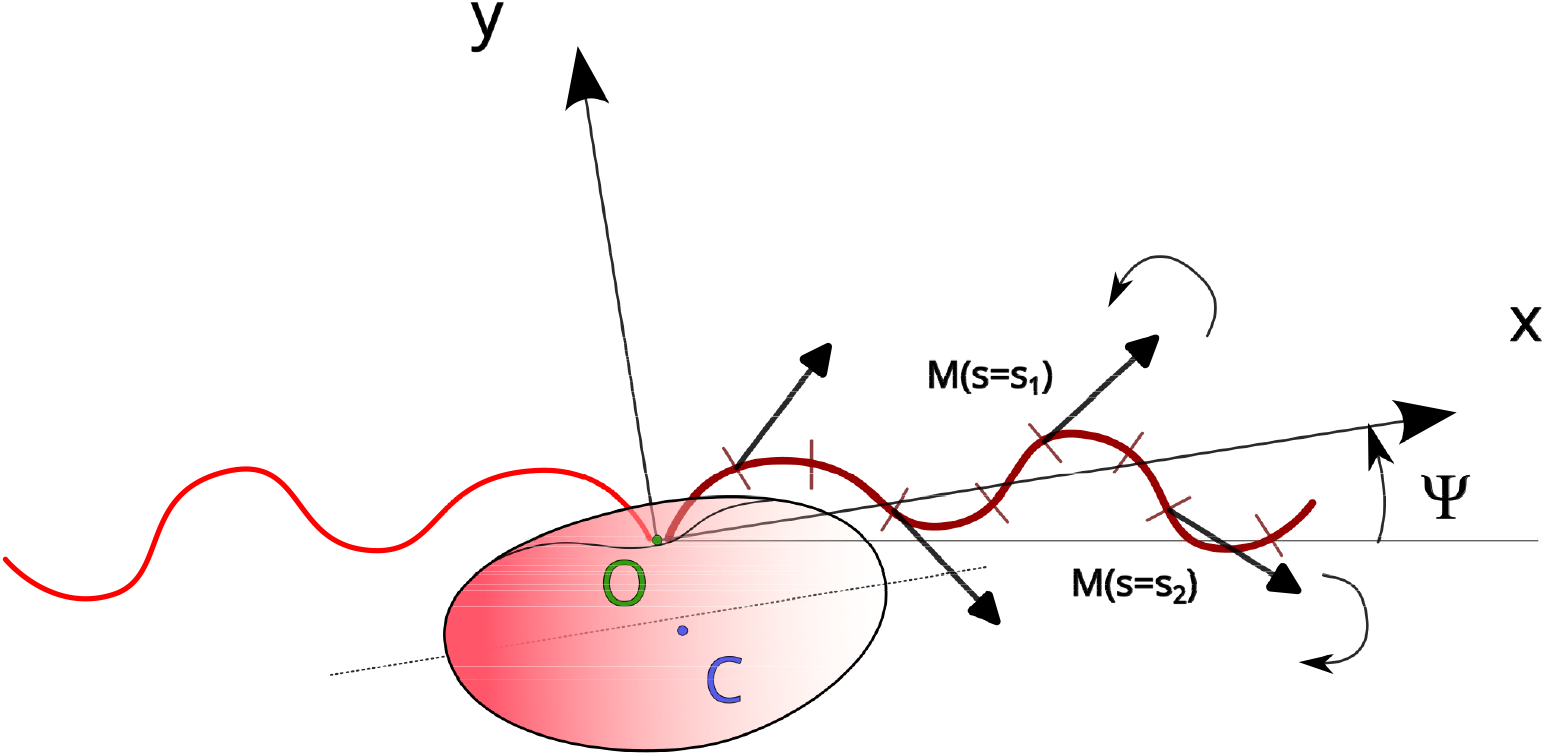
Zoospores, flagella with forces and definition of angle Ψ.

## 2 Perturbative analysis

In this first step, we present a way to determine the angular speed reached by the cell body (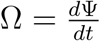. This quantity is directly linked to the amplitude of the angular oscillation. We assume that a perturbative analysis can be performed, with velocity components linked only to the translation along x axis, being **of** order 0, and with a rotation being of order 1. Only the order 0 will be used to estimate the value of Ψ(t).

In a second step, the rotational speed can be introduced, leading to coupled equations for 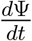.

Following the analysis of [2] The force written : 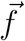 by unit length along the flagella can bex

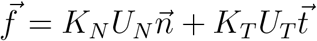

It has been shown at 4 different locations on the flagella in Fig.1. The local force in s = s_1_ tends to rotate counter-clockwise the cell body (cb), and the contribution from force in s = s_2_ clockwise. We see here that depending on the phase of the flagella beating, the different contributions will compensate except the last **one**, which will present a positive or negative sign inducing a rotation in a given direction. We present here the calculation for one **flagellum**, knowing that the same applies for the other. **Since the** Stokes equation is linear, the total rotation induced to the cell body will be the sum of both contributions from **the** anterior and posterior flagella.

### 2.1 Force balance

The force balance over the total length of a flagella can be written **as follows** :

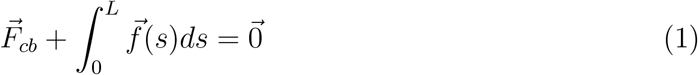

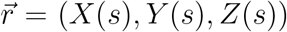, is the position vector of the point M(s) along the flagella for 0 < s < L. Z(s) = 0 for the moment for simplicity. 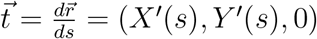

The velocity u is given in the standard theory with no rotation of the cell body by (we keep only order 0) :

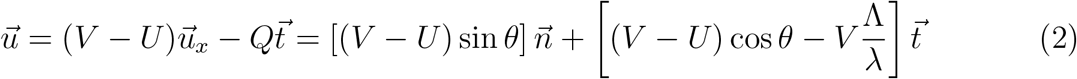

So :

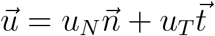

sθ is the angle between 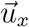 and 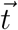. Q is the wave speed along the flagellum, it corresponds to a wave speed V along x axis : 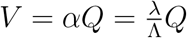. 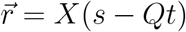,Y (s − Qt), 0).

The force 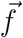 is then equal to :

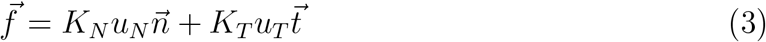

The total force on the cell body is then given by :

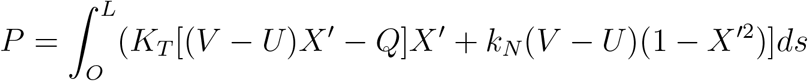

Noting 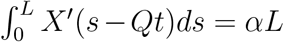 and 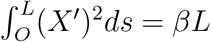, and since V = αQ we then have :

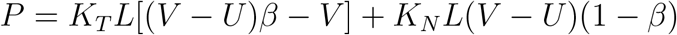

Taking only the drag from the head leads to : P = K_*N*_ Lδ. By equating the forces, noting ϕ = K_*T*_ /K_*N*_, we obtain the classical result of RFT (see [2], Eq. 7.16) :

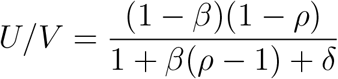

### 2.2 Torque balance

For the torque balance, as taken along the z axis passing in C :

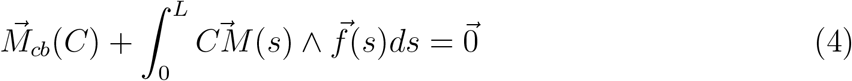

which can be rewritten :

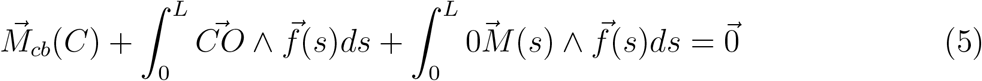

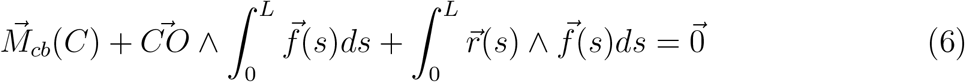

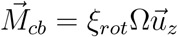. Ω being the instantaneous angular speed of the cell body.

We need now to assume that the cell body can have his angle **compared** to fixed u_*x*_ axis in the lab reference frame, oscillating, in a range inferior to θ, we note it Ψ. We have : 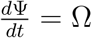

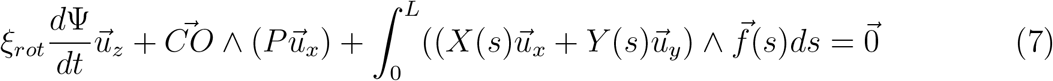

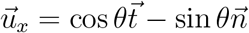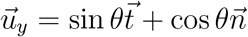

X^I^(s − Qt) = cos θ.

Q = Λν, the wave speed along the flagella.

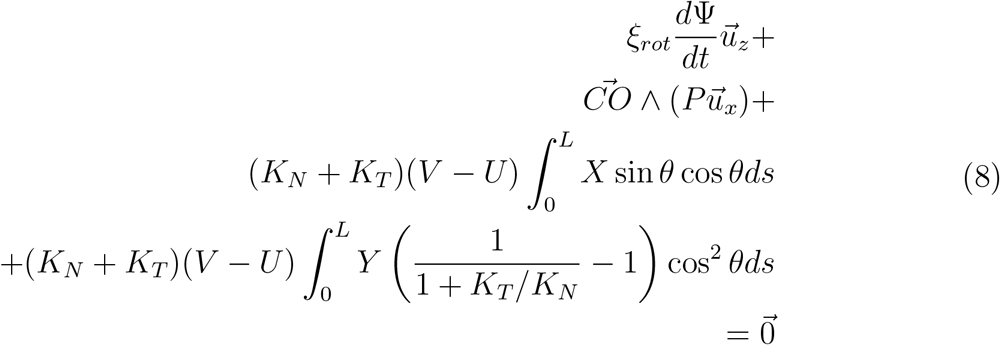

To calculate the two integrals I_1_, I_2_ and have an estimate for the evolution of the angle, we need to calculate :

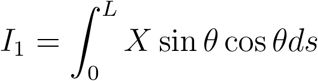

and

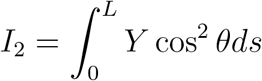

Knowing that cos 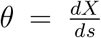. Either the shape as function of time is known (X(s − Qt),Y (s − Qt)). For example, we could assume : 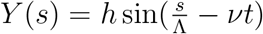 and do numerical analysis of the problem.

To perform an analytical calculation, we first can move to a description in term of x variable instead of s. We assume 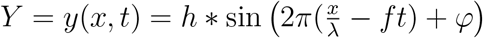.

Then 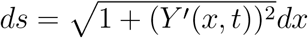.

We have 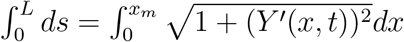, and in particular 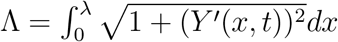.

We have then :

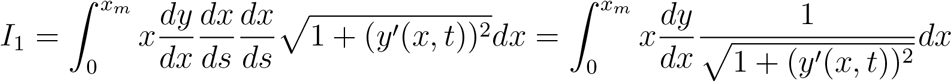

by noting 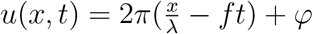, we have then :

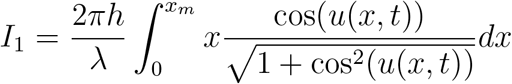

and following the same idea,

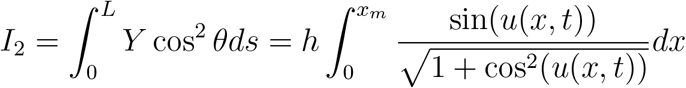

To go beyond, we can assume small amplitude for the flagella (h « λ, leading to :

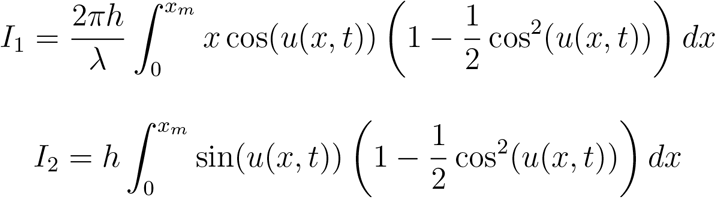

Both these integrals can be calculated by parts and lead to analytical expressions with various terms of the form 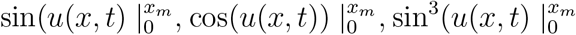 and 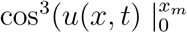.

This lead to sum of functions of the form : cos(2πf t+ϕ_1_), cos(2πft+ϕ_2_), cos(2π(3ft)+ϕ_3_), cos(2π(3ft)+ ϕ_4_).

In the end, using Eq.8, performing the same analysis with the third integral gives at leading order constant values, and only cubic terms.

The analysis has to be performed similarly for the two flagella and both contributions have to be added.

In the end, assuming this perturbation analysis and small deformation for the flagella, we end up with the function Ψ(t) below (Eq. 9. It contains a linearly increasing term linked to constant rotation rate (a_0_) if the effects of the two flagella do not cancel, plus a term at beating frequencies of both flagella, plus terms of harmonics at 3 times the frequencies.

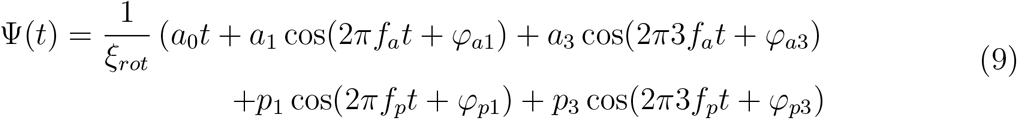

The amplitude of oscillations of the cell body angle are directly proportional to the flagellar beating amplitude, inversely proportional to ξ_*rot*_ and can depends strongly on the ratio of the length of the flagella to the wavelength which have a strong impact on the coefficient a_1_, a_3_, p_2_, p_3_. This parameter is the most important and depending of its value will make positive, null or negative the oscillation angle of the cell body.

Another complexity in the problem is the fact that the cell body gyrates on itself, as described in [1]. This implies an additional effect not taken into account here which is a torque along the main direction of motion, and not perpendicular to the beating plane as studied here. The main effect of this gyration is to constantly turning the plane of flagellar beating. when the plane is perpendicular to the optical axis of the camera, the beating amplitude is the largest, but when the plane contains the optical axis, the beating amplitude and the cell body oscillation decreases significantly, as seen on the Fig. 1 of the main text of the manuscript.

To conclude, we show here with approximations of small deformation and assuming small perturbation from the angular oscillation of the cell body that the Fourier spectrum of the cell body angle variations is made of two main peaks at the frequency of beating of both anterior and posterior flagella. Then smaller peaks at 3 times the beating frequencies of each flagella should be present. Finally a more or less broad peak close to null frequency linked to the rotation with coefficient a_*O*_ in Eq. 9 which is slowly variable with time. It characterizes the more or less rapid turning events and curved trajectories. The Fourier mode amplitudes at beating frequency are an average value over a whole gyration on itself of the cell body, underestimating by a factor of about 2 its real amplitude.

## 3 Full coupled model

A more complex and exact model would require numerical analysis to limit the approximation on small deformations, to write torque equation along the propulsion axis to take into account the gyration on itself of the cell body, and to add the rotation term in the velocity kinematic equation (Eq.2). This latter equation is modified since the velocity components are now written fully, and presents more terms :

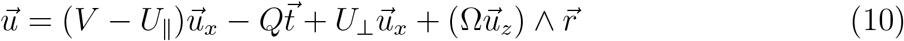

With 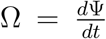. Then Eq. 7 has to be written with Eq. 3 with this new **relation**, projected on 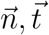.

In this new equation, the two terms with 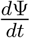 need to be assembled. The full determination is out of the scope of the present analysis which was mainly focused on explaining the characteristics of the Fourier spectrum measured.

